# Smallpox vaccination induces a substantial increase in commensal skin bacteria that promote pathology and enhance immunity

**DOI:** 10.1101/2021.07.30.454438

**Authors:** Evgeniya V. Shmeleva, Mercedes Gomez de Agüero, Josef Wagner, Anton J. Enright, Andrew J. Macpherson, Brian J. Ferguson, Geoffrey L. Smith

## Abstract

Interactions between pathogens, host microbiota and the immune system influence many physiological and pathological processes. In the 20^th^ century, widespread dermal vaccination with vaccinia virus (VACV) led to the eradication of smallpox but how VACV interacts with the microbiota and whether this influences the efficacy of vaccination are largely unknown. Here we report that intradermal vaccination with VACV induces a large increase in the number of commensal bacteria in infected tissue, which enhance recruitment of inflammatory cells, promote tissue damage and increase immunity. Treatment of vaccinated specific-pathogen-free (SPF) mice with antibiotic, or infection of genetically-matched germ-free (GF) animals caused smaller lesions without alteration in virus titre. Tissue damage correlated with enhanced neutrophil and T cell infiltration and levels of pro-inflammatory tissue cytokines and chemokines. One month after vaccination, GF mice had reduced VACV-neutralising antibodies compared to SPF mice; while numbers of VACV-specific CD8^+^ T cells were equal in all groups of animals. Thus, skin microbiota may provide an adjuvant-like stimulus during vaccination with VACV. This observation has implications for dermal vaccination with live vaccines.

**Author Summary:** Smallpox was caused by variola virus and was eradicated by widespread dermal vaccination with vaccinia virus (VACV), a related orthopoxvirus of unknown origin. Eradication was declared in 1980 without an understanding of the immunological correlates of protection, or knowledge of the effect of smallpox vaccination on the local microbiota. Here we demonstrate that intradermal infection of mice with VACV induces a ∼1000-fold expansion of commensal skin bacteria that influence the recruitment of inflammatory cells into the infected tissue and enhance the size of the vaccination lesion. Antibiotic treatment reduced lesion size without changing virus titres. The bacterial expansion also contributes to the level of neutralizing antibodies at one month post vaccination, because genetically matched germ-free mice developed lower neutralizing antibodies than specific pathogen free controls. Thus, dermal infection by VACV enhanced bacterial growth and these bacteria promote pathology and enhance the antibody response. This finding has implication for dermal vaccination with live vaccines.

## Introduction

Vaccinia virus (VACV) is the live vaccine that was used to eradicate smallpox, a disease caused by the related orthopoxvirus, variola virus, and declared eradicated by the World Health Organisation in 1980 (1). During the smallpox eradication campaign, early vaccine batches were grown on the flanks of animals and, consequently, were not bacteriologically sterile. Vaccination was achieved by multiple skin puncture using a bifurcated needed containing a drop of vaccine suspended between the two prongs of the needle. Therefore, both the vaccine and the method of vaccination may have introduced bacteria into the vaccination site. Following vaccination a local lesion developed that healed in 2-3 weeks and left a characteristic vaccination scar. A careful study of smallpox vaccination outcomes in the USA described several serious dermal or neurological complications of vaccination (2), although bacterial infection of the vaccination site was not highlighted. Vaccination induced long-lasting humoral and cellular responses, although the precise immunological correlates of vaccination remain uncertain (3). VACV encodes scores of proteins that block the innate immune response and induce local immunosuppression (4) and manipulation of these proteins can affect virulence and adaptive immunity (5).

Secondary bacterial infection is sometimes a consequence of viral infection; classic examples are bacterial pneumonia following influenza virus, respiratory syncytial virus or parainfluenza virus infection (6). These bacterial infections may be promoted by virus-induced tissue damage facilitating bacterial entry into tissue, loss of anti-microbial proteins, or virus-induced immunosuppression enabling invasion by commensal organisms. Bacteria causing these secondary infections are often derived from the host microbiota (7, 8). Bacterial infection has been described following infection by several poxviruses including avipoxvirus (9), cowpoxvirus (10), monkeypox virus (11) and variola virus (12) although this was not highlighted as a complication of smallpox vaccination (2).

Vaccine development can be a slow and expensive process and understanding how to deliver antigens optimally to induce strong adaptive immunity remains poor. In particular, understanding how innate immunity leads to development of adaptive immunity is incomplete (13, 14). The activation of dendritic cells by innate immunity enhances antigen presentation and thereby improves the adaptive memory response (15), but innate immunity may also cause swift elimination of antigens, and so diminish adaptive immunity (16). Since bacterial components are sometimes used as adjuvants to enhance innate immunity and vaccine efficacy (17), studying the role of microbiota following vaccination is important.

In this study, the mouse intradermal model of VACV infection (18) was used to make a detailed investigation of the recruitment of inflammatory cells into the infected tissue. Previously, the cellular infiltrate in this model was found to contain abundant neutrophils and T cells and was markedly different to the response to respiratory infection (19). Neutrophil recruitment is a hallmark of bacterial infection (20) and this led to the discovery here that the local skin microbiota expanded greatly following dermal VACV infection. Data from vaccinated antibiotic-treated or germ free (GF) mice showed that the pathology at the site of the viral infection is dominated by commensal bacteria, without influencing virus replication. Notably, depletion of bacteria resulted in alteration to both the cellular infiltration into infected tissue and the subsequent adaptive immune response.

This study highlights a role for commensal bacteria in enhancing the immune response following dermal vaccination and has implication for other vaccines based upon infectious poxviruses or other viral vectors that are delivered by dermal vaccination.

## Results

### Neutrophils infiltrate ear tissue after intradermal vaccination with vaccinia virus

Intradermal (i.d.) infection of the mouse ear pinnae with VACV strain Western Reserve (WR) results in the development of a skin lesion at the site of infection that mimics smallpox vaccination in man. A lesion appears at d 6 post infection (p.i.), increases gradually to about d 10 and resolves by d 21 (18). The model has been used to study the contribution of individual VACV proteins to virulence and immunogenicity (21, 22).

Here the local innate and cellular immune responses to i.d. vaccination with VACV were characterised in detail, including quantifying subpopulations of infiltrating myeloid cells throughout infection. Flow cytometry showed a substantial increase in CD45^+^CD3^-^CD5^-^CD19^-^NK1.1^-^CD11c^-^Siglec-F^-^Ly6G^+^ cells (neutrophils) in ear tissue at all times p.i. (Fig 1A, B). Along with the surface markers, the morphological characteristics of this population (spherical ∼8-10 µm cells with highly segmented nuclei) confirmed their identity as neutrophils (S1 Fig) (23). The number of neutrophils increased up to 10 d p.i., and by d 5 had increased ∼130 fold compared to mock-infected tissue (Fig 1B), coinciding with the first appearance of a dermal lesion. Neutrophils are part of the innate immune system and are front-line effector cells for defense against bacterial infection (20). Their abundance in VACV-infected tissue suggested that bacterial co-infection might be present.

**Fig 1.**
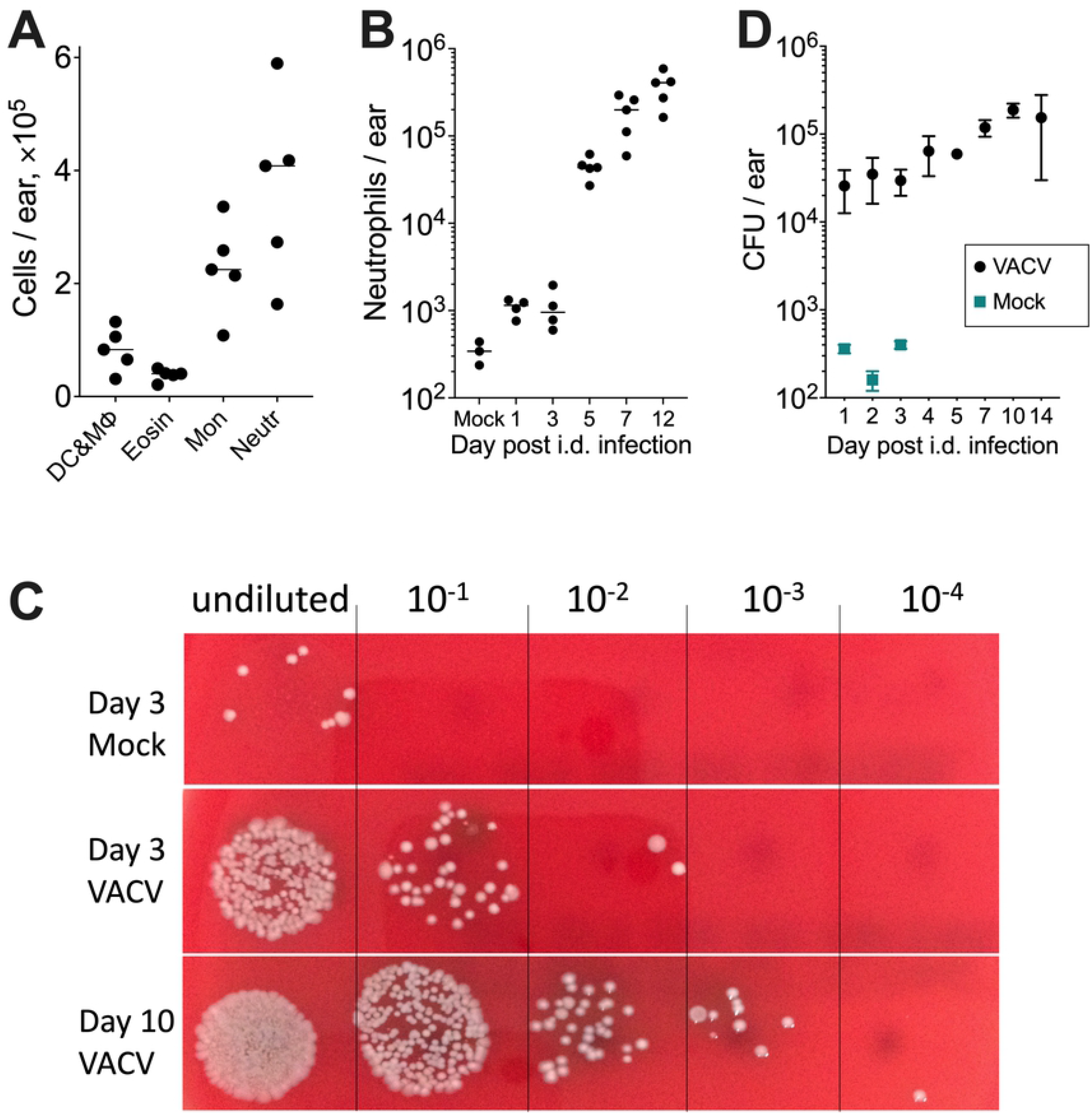
Skin microbiota expand after VACV infection. C57BL/6 SPF mice were injected i.d. with 10^4^ PFU of VACV strain WR or PBS (mock) and ear tissues were then collected at different times post injection. (A) Absolute numbers of different myeloid cells present in ear tissues at d 9 p.i. (*n*=5 per time point). DC & MФ: dendritic cells and macrophages; ; Eosin: eosinophils; Mon: monocytes; Neutr: neutrophils. Medians are shown. (B) Absolute numbers of neutrophils infiltrating ear tissues at different times p.i. Mock: mock-control, 5 d post intradermal injection of PBS (*n*=3-5 per time point). Medians are shown. (C) Bacterial colonies grown from homogenised ear tissues and their 10-fold serial dilutions seeded on blood agar. (D) Bacteria colony-forming unit (CFU) counts of ear samples (n=3). Means and SEM are shown. The experiments were performed at least twice and representative data from one experiment are shown.

### Skin microbiota expand after VACV infection

To investigate if there is an expansion of bacteria after i.d. infection with VACV, bacterial colony-forming units (CFUs) in infected tissue were quantified. CFUs in VACV-infected tissue was at least 100-fold greater than mock-infected tissue at all time points (Fig 1C) and at the peak at 10 d p.i. was about 1000-fold greater than control (Fig 1D). Given that the virus inoculum was bacteriologically sterile, these results suggest increased growth of commensal bacteria after VACV infection, possibly due to local virus-induced immunosuppression. Notably, the time of maximum number of bacteria and neutrophils correlated with maximum lesion size, suggesting a link between skin microbiota and the severity of pathology.

### Skin microbiota composition changes after VACV infection

To investigate the identity of the bacteria present during infection and to compare this with the microbiome in uninfected skin, 16S ribosomal RNA gene (rRNA) sequencing was performed. So that bacteria from the skin surface and within lesions was included, total genomic DNA was extracted directly from whole VACV-infected and non-infected ear tissues and next generation sequencing of the amplified V4 region of bacterial 16S rRNA was performed. Sequencing was carried out from material extracted directly from infected tissues without prior culturing to avoid bias introduced by the culturability of bacteria that might be present.

Almost all the bacterial families found belonged to: 1) *Firmicutes* (*Rhodobacteraceae, Lachnospiraceae, Planococcaceae, Aerococcaceae, Clostridiaceae, Carnobacteriaceae, Bacillaceae, Staphylococcaceae, Lactobacillaceae, Streptococcaceae, Clostridiaceae, Ruminococcaceae, Veillonellaceae, Erysipelotrichaceae*); 2) Actinobacteria (*Micrococcaceae, Corynebacteriaceae, Actinomycetaceae, Dietziaceae, Nocardiaceae, Propionibacteriaceae, Nocardioidaceae, Microbacteriaceae*); 3) *Proteobacteria* (*Aurantimonadaceae, Methylobacteriaceae, Sphingomonadaceae, Burkholderiaceae, Comamonadaceae, Oxalobacteraceae, Neisseriaceae, Desulfovibrionaceae, Alteromonadaceae, Enterobacteriaceae, Halomonadaceae, Pasteurellaceae, Moraxellaceae, Pseudomonadaceae, Caulobacteraceae, Xanthomonadaceae*); and 4) Bacteroidetes (*Bacteroidaceae, Porphyromonadaceae, Prevotellaceae, Chitinophagaceae, Cytophagaceae, Flavobacteriaceae*) (S2A Fig). These are consistent with mouse skin microbiota reported previously (24, 25), suggesting that the source of the enhanced numbers of bacteria in VACV-infected tissue is the skin rather than environment.

Analysis of bacterial sequences at the family, genus and species levels showed significant changes in the composition of skin microbiota following infection with VACV (Fig 2, S2 and S3 Figs). Although mock-infection itself resulted in moderate modification of skin microbiome, the most significant changes occurred at d 5, 8 and 12 p.i., especially d 8 and 12, which was clearly demonstrated by principal component analysis (Fig 2A, S2B and S3A Figs) as well as multivariate beta diversity analysis (S1 Table). The relative abundance of some dominant genera in intact skin such as *Sporosarcina* and *Staphylococcus* decreased at late times post VACV infection, while the proportion of *Streptococcus, Enhydrobacter* and *Corynebacterium* increased (Fig 2B). Perturbation of skin microbiome also occurred within the same genus, for instance VACV infection resulted in a rise of relative abundance of *Staphylococcus aureus*, a well-known opportunist, over other species of *Staphylococcus* (S3B Fig). While averaged data visualised as stack plots cannot represent in full the observed microbiota shift, heatmaps (Fig 2C, S2A and S3C Figs) illustrate that the majority of d 8 and 12 VACV-infected samples had very pronounced dominance of ∼1-3 bacterial genera, which differs from normal microbiota. However, specific dominant taxa varied from sample to sample. Therefore, there is no particular type of bacterium that spread favourably after VACV infection. Instead, random different bacteria are increased in infected samples. Thus, VACV infection leads to changes in the skin microbiome and the expansion of opportunistic bacteria.

**Fig 2.**
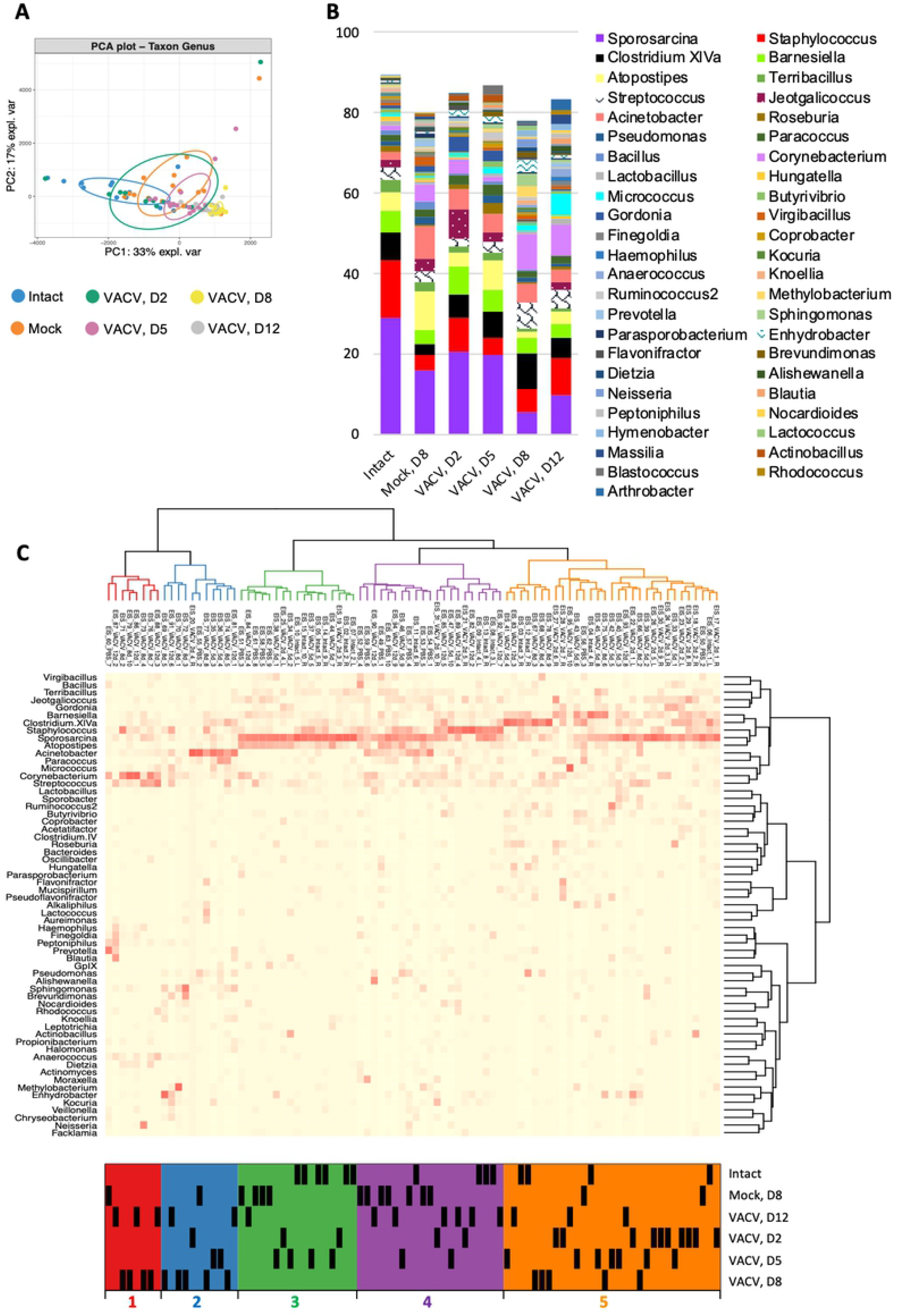
Skin microbiome change after i.d. infection with VACV. Ear tissues were collected from SPF mice before (intact) and at 2, 5, 8 and 12 d p.i. with 10^4^ PFU of VACV and at d 8 post PBS injection (mock), *n* = 15 per group per time point. Next generation sequencing was performed for DNA samples extracted from tissue. (A) Principal Component Analysis (PCA) of microbiome of genus taxon. PCA was conducted including all taxon with a minimum abundance of 0.1%. Ellipses represent a 40% confidence interval around the cluster centroid. (B) Relative abundance of most prevalent taxa of bacterial genera. (C) Heatmap of bacterial genera with hierarchical clustering. Bacterial genera with a minimum abundance of 1% in at least five samples was used for creation of the heat map. Five clusters are colour-coded with red, blue, green, violet and orange. “D” – day post injection.

### Skin microbiota promote lesion development after VACV infection

To investigate the influence of skin microbiota on lesion formation, a broad-spectrum bactericidal antibiotic (AB, ceftriaxone) was administered intraperitoneally (i.p.) over 13 d after VACV infection to suppress bacterial growth. C57BL/6 specific-pathogen-free (SPF) mice were infected i.d. with VACV, and one group received antibiotic treatment (SPF, AB-group), while another received phosphate-buffered saline (SPF, NoAB-group). Antibiotic treatment caused a significant reduction in lesion size after infection with VACV strain WR, a widely used laboratory strain of VACV (Fig 3A). A similar observation was made following infection with VACV strain Lister, a strain used widely for smallpox vaccination in humans (S4 Fig). To confirm that this effect was due to the presence of bacteria, rather than a non-specific consequence of antibiotic treatment, genetically-matched germ-free (GF) mice were infected i.d. with VACV strain WR. These mice showed delayed lesion formation and had a three-fold reduction in lesion size compared to SPF animals (Fig 3B).

**Fig 3.**
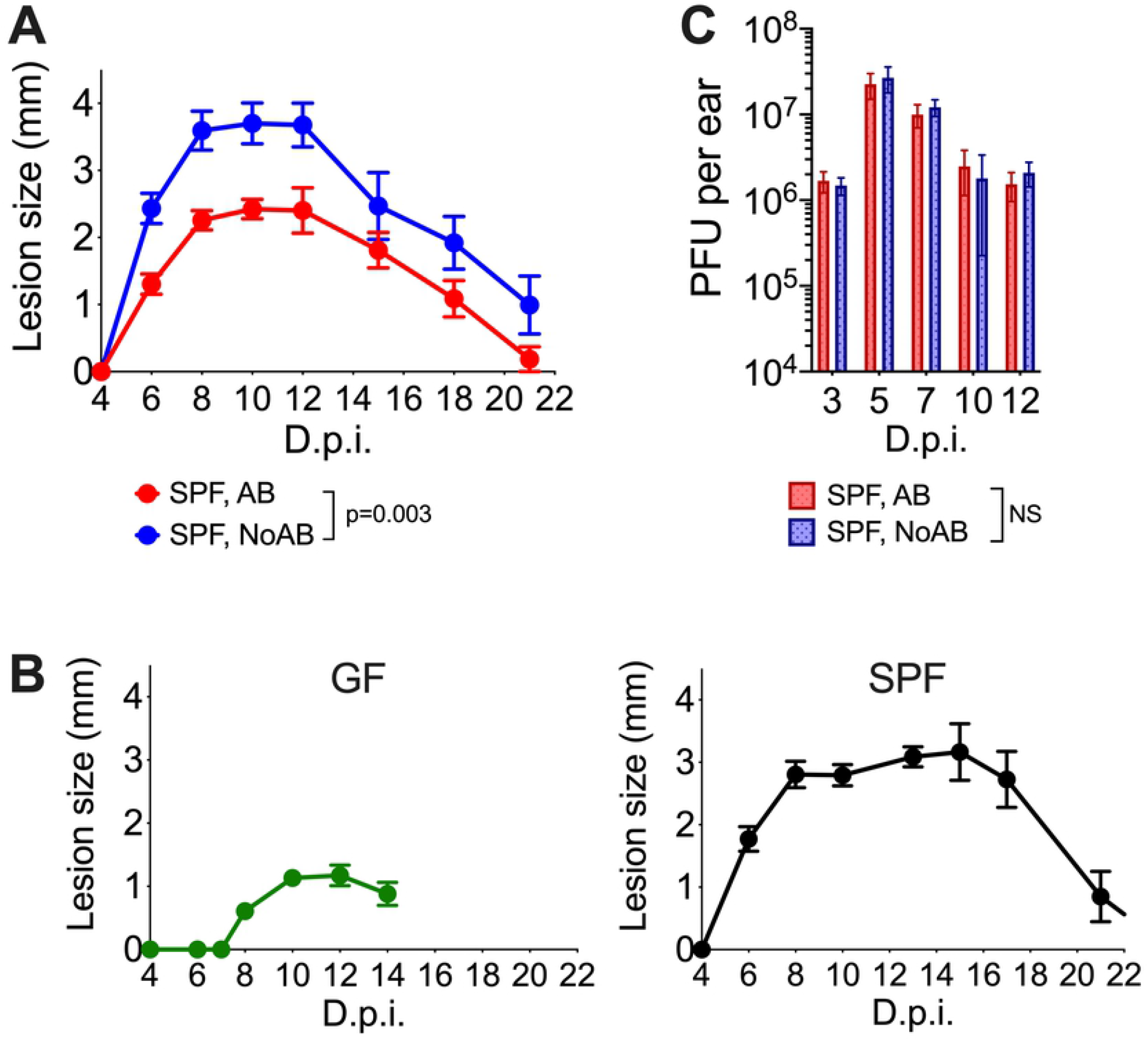
Skin microbiota promote lesion development after VACV infection. Specific pathogen free (SPF) or germ-free (GF) mice (n=5-15 per group) were injected intradermally with 10^4^ PFU of VACV. SPF, AB animals received antibiotic i.p. from 1 d p.i.; SPF, NoAB animals received i.p. injections with PBS. (A-B) Ear lesion sizes of AB and NoAB-groups (A) or GF or SPF mice (B). Means and SEM are shown. Statistical analysis by two-way RM ANOVA test. (C) VACV titres in ear tissue p.i. PFU, plaque-forming units. NS – non-significant by Mann-Whitney test. Experiments (except for those with GF mice) were performed at least twice and representative data from one experiment are shown. The experiments with GF mice were performed once.

Topical administration of antibiotic creams on the infected ears also resulted in a reduction of lesion size following VACV infection (S5 Fig). However, it was not possible to control the dose administered because the animals tend to groom themselves and each other, leading to ingestion of the applied creams. Antibiotic treatment can influence antiviral immunity when administered orally (26) and so to minimise the influence on gut microbiota, the antibiotic was administered i.p. for all our experiments. To avoid changes in microbiota prior to infection, antibiotic treatment was only started from d 1 p.i. Throughout ceftriaxone treatment, the health status of animals and their leukocyte composition in peripheral blood, spleen and bone marrow were unaltered (S6 Fig).

Next the virus titres in infected tissue with or without antibiotic treatment were determined. Notably, despite considerable differences in lesion size, the viral titres in infected ear tissues were unaltered (Fig 3C). This indicated that: i) antibiotic treatment has no impact on virus replication, and ii) lesion size was influenced by the presence of bacteria rather than virus titre.

Histological examination of infected lesions showed that necrosis of the ear tissue in the NoAB-group of animals was substantially greater than in the AB-group (Fig 4A). A striking difference in histological changes after VACV infection was also observed when comparing GF and SPF mice: ears of the GF animals had significantly reduced cell infiltration and only very mild tissue necrosis (Fig 4B).

**Fig 4.**
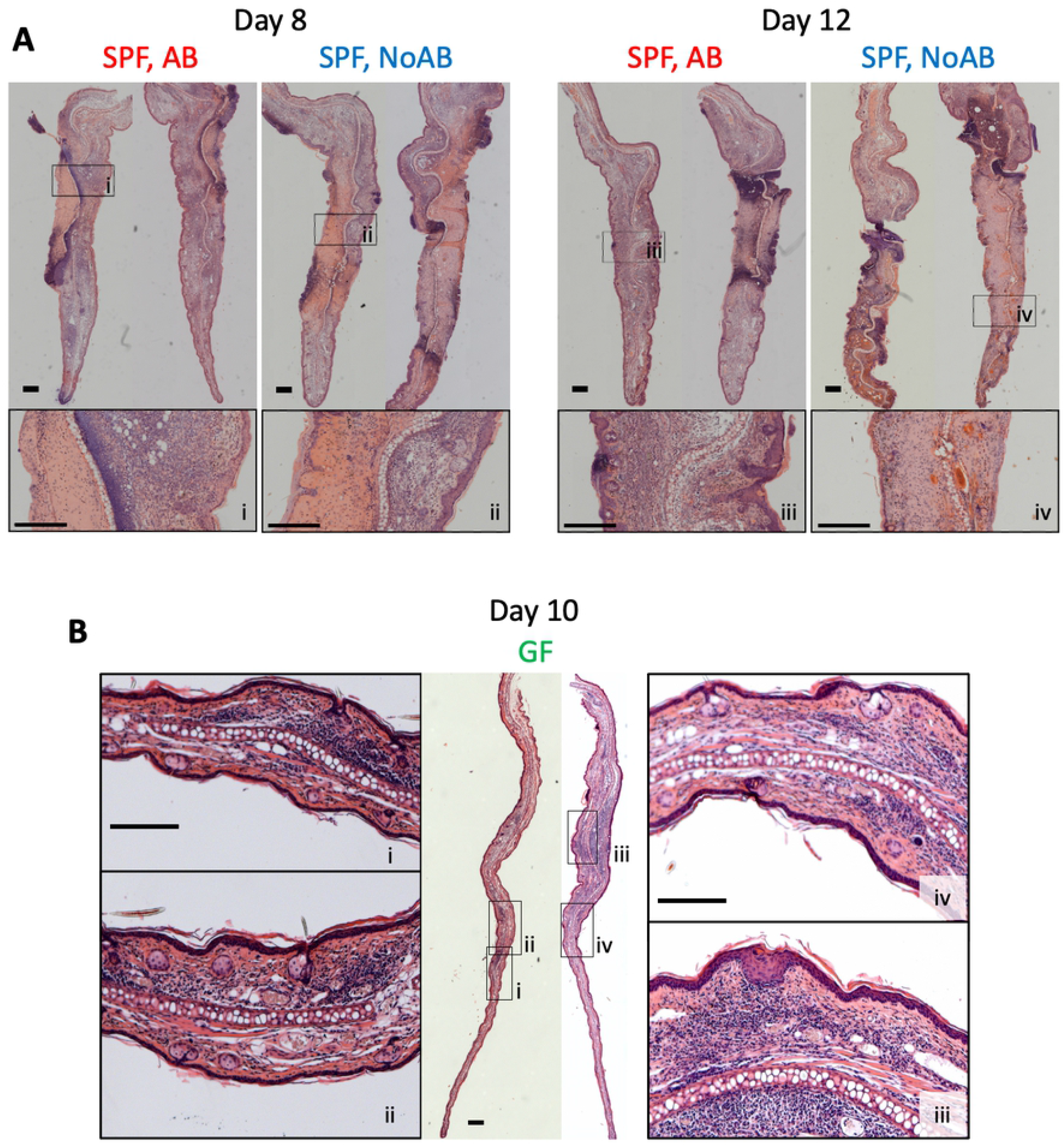
Histology of ear lesions after VACV infection. Specific pathogen free (SPF) or germ-free (GF) mice (*n*=5-15 per group) were injected i.d. with 10^4^ PFU of VACV WR. Group SPF AB was injected daily i.p. with antibiotic ceftriaxone from d 1 to 11 p.i. Group SPF NoAB received i.p. injections with PBS. (A) Images of haematoxylin and eosin (H&E) stained transverse ear sections 8 and 12 d p.i. Bars = 300 µm. (B) Images of H&E stained transverse ear sections collected from GF mice 10 d p.i. Bars = 300 µm.

Collectively, these observations indicate that skin microbiota promote lesion development after VACV infection and the smaller lesion sizes and lack of pathological tissue changes in GF and antibiotic-treated groups were due to the absence of microbiota or antibiotic-induced suppression of bacterial expansion.

### Skin microbiota enhance immune cell recruitment and cytokine production

Next, we compared the local immune response to VACV infection in ear tissue of antibiotic-treated and untreated SPF animals. Recruitment of immune cells was low on d 1-4 p.i. (Fig 5A) but there was a sharp increase of Ly6C^+^ inflammatory monocytes on d 5, followed by large leukocyte infiltration at d 7. Notably, the antibiotic-treated group showed a substantial reduction in the recruitment of multiple subpopulations of myeloid and lymphoid cells at d 7 p.i. (Fig 5A). By d 12 p.i., several subpopulations of leukocytes had declined, except for neutrophils in the NoAB-group, which continued to increase. During infection there was a notable positive correlation between the lesion size and the numbers of infiltrating neutrophils or TCRαβ T cells and this correlation did not hold for any other cell type (Fig 5B).

**Fig 5.**
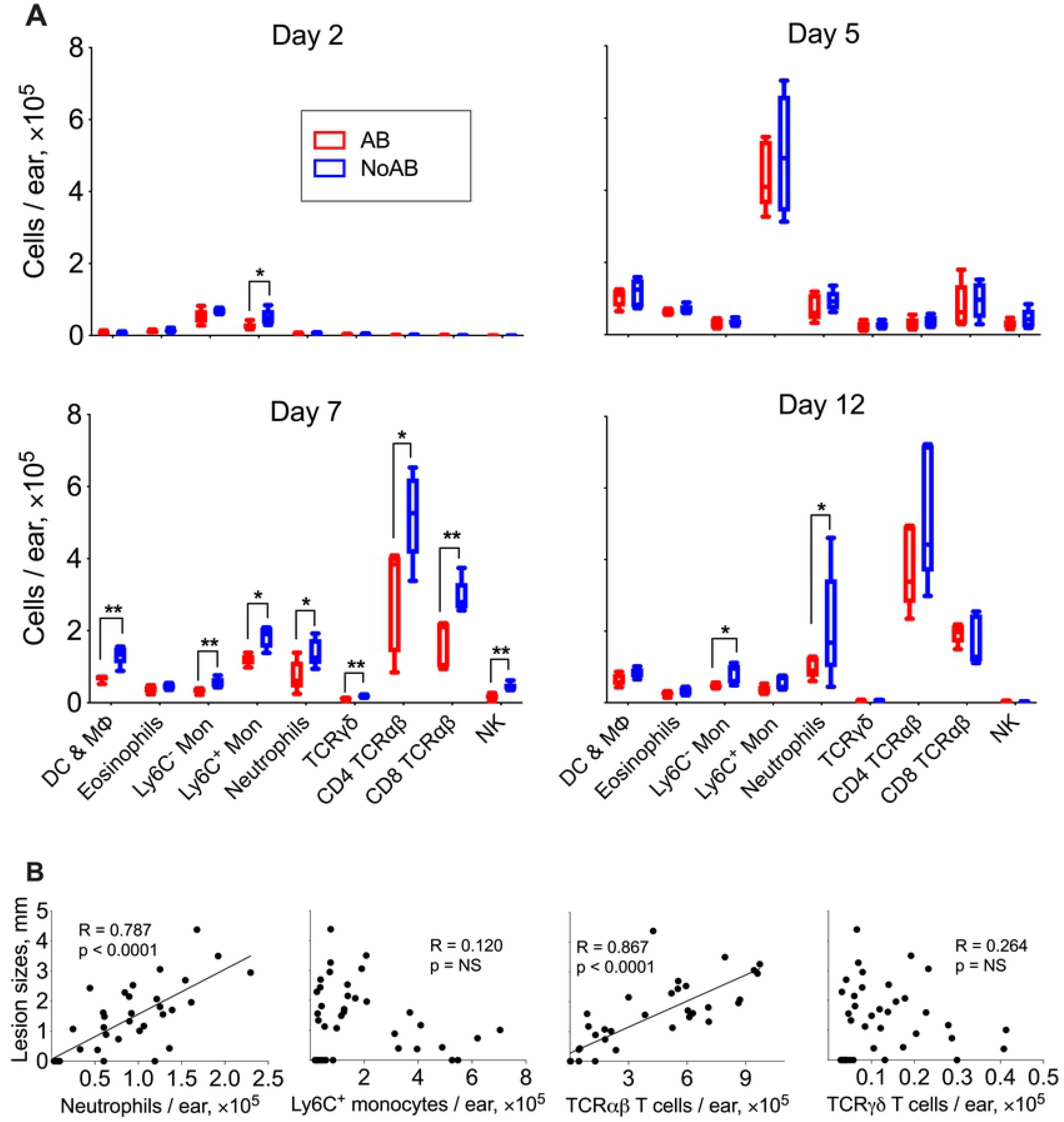
Skin microbiota advance immune cell recruitment into VACV infected tissue. Two groups of SPF mice were infected i.d. with 10^4^ PFU of VACV. From d 1 p.i. onwards, one group (AB) received antibiotic (ceftriaxone) i.p. daily. The second group (NoAB) received PBS i.p. (A) Numbers of myeloid and lymphoid cell of subpopulations were measured in ear tissues at 2, 5, 7 and 12 d p.i. (n=5 per group per time point). DC & MФ, dendritic cells and macrophages; Mon, monocytes, TCRγδ, TCRγδ^+^ T cells; TCRαβ CD4, TCRγδ^-^CD4^+^ T cells; TCRαβ CD8, TCRγδ^-^CD8^+^ T cells. Box plots are shown; p values were determined by the Mann-Whitney test, * = p<0.05, ** = p<0.01. (B) Spearman correlation analysis of lesion sizes versus numbers of neutrophils, Ly6C+monocytes, TCRγδ T cells or TCRαβ (TCRγδ^-^) T cells recruited to site of infection. The experiment was performed twice and representative data from one experiment are shown.

To analyse the local inflammatory environment further, we measured an array of cytokines and chemokines in the ear tissue at d 5, 8 and 12 p.i. in control and antibiotic-treated mice. Consistent with the smaller lesions and reduction in cellular infiltration, among 17 different inflammatory mediators measured, there were reductions of IFNγ, TNFα, CCL2, CCL4, CCL7, CXCL1 and CXCL10 levels in the antibiotic-treated group compared to controls at d 8, and also in the amount of CCL4 and CCL7 at d 12 p.i. (Fig 6A). Similar differences were also evident when comparing infected GF mice with SPF mice. Levels of IFNγ, TNFα, CCL2, CCL7, CXCL1 and CXCL10 were considerably diminished in the GF mice (Fig 6B). These data indicate that the presence of skin microbiota accelerates overall immune cell recruitment and cytokine/chemokine production in VACV-infected skin.

**Fig 6.**
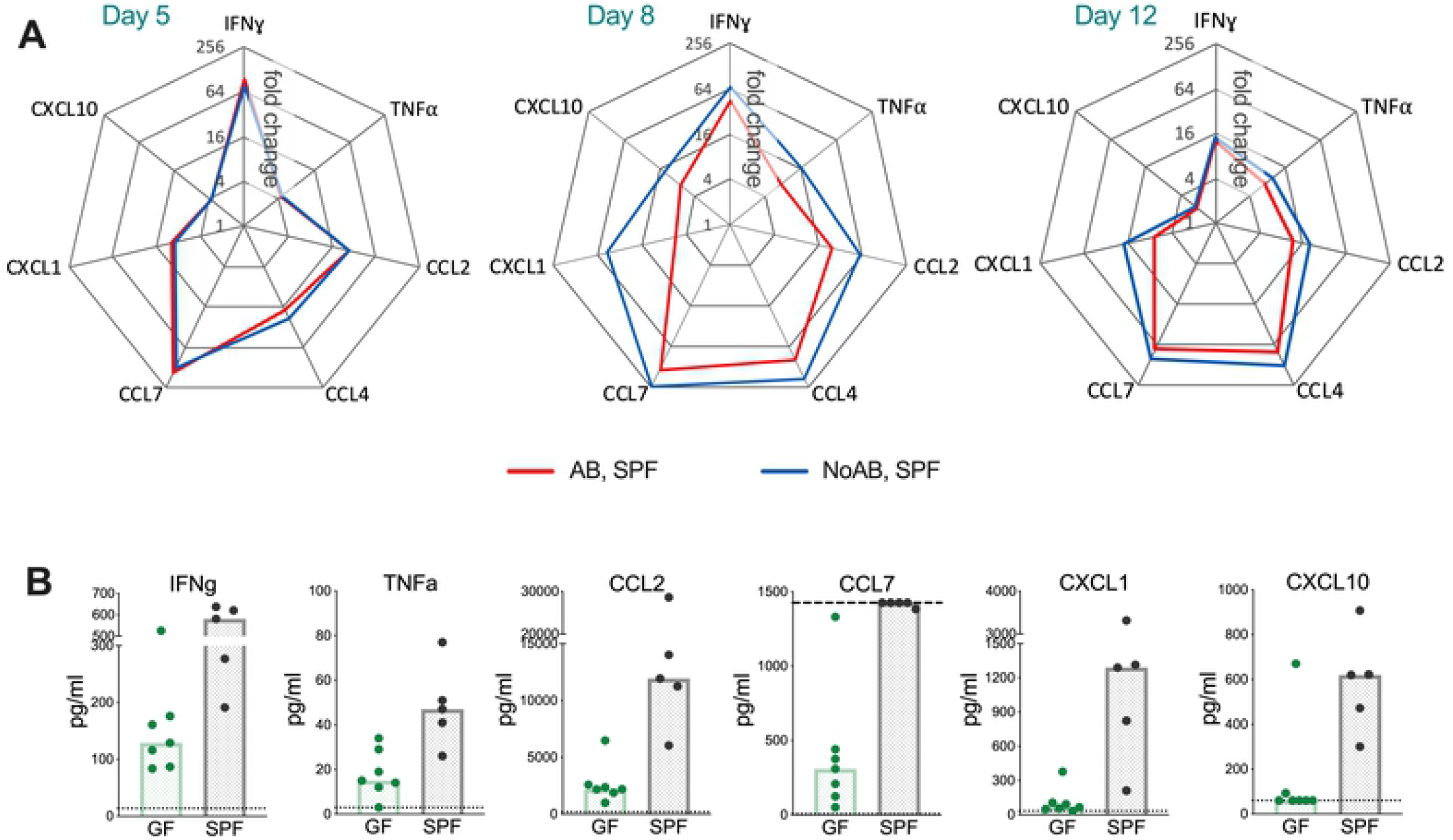
Skin microbiota promote the production of cytokines/chemokines in VACV-infected ear tissue. SPF or GF mice (n=5-7 per group per time point) were injected i.d. with 10^4^ PFU of VACV. SPF, AB-group received antibiotic (ceftriaxone) i.p. daily from 1 d p.i.; SPF, NoAB group received i.p. injections with PBS. (A) Tissue levels of cytokines and chemokines were measured by multiplex assay (Luminex). Data are shown as the fold change from the baseline (untreated intact ear samples). Means are shown. The experiment was performed twice and data from one representative experiment are shown. (B) Tissue levels of cytokines and chemokines in GF and SPF groups at d 10 p.i. measured by multiplex assay (Luminex). Graphs show results two independent experiments. Dotted lines indicate the lowest standards (or highest standard for CCL7). The experiment with GF animals was performed once.

### Humoral adaptive immune response is impaired in the absence of microbiota

To assess the role of microbiota in the generation of VACV-specific adaptive immunity, we measured VACV-specific CD8^+^ T cell numbers and anti-VACV neutralising antibodies in samples obtained from SPF groups, with or without antibiotic treatment, and GF mice one month after vaccination. Absolute numbers of splenic CD4^+^ and CD8^+^ T cells, as well as VACV-specific CD8^+^ cells, were similar in all three groups of animals (Fig 7A, B). However, the levels of VACV neutralising antibodies were significantly lower in GF animals than in either group of SPF animals (Fig 7C). Despite the reduced antibody response in the vaccinated GF mice, all groups showed protection from disease upon intranasal challenge with a lethal dose of VACV (Fig 7D-F). Each SPF group showed a small (∼10%) transient weight loss after challenge before recovery. These data indicate that the GF mice have an impaired capacity to generate neutralising antibody following i.d. vaccination with VACV, but that this does not affect their capacity to develop protective immunity against re-challenge in this model.

**Fig 7.**
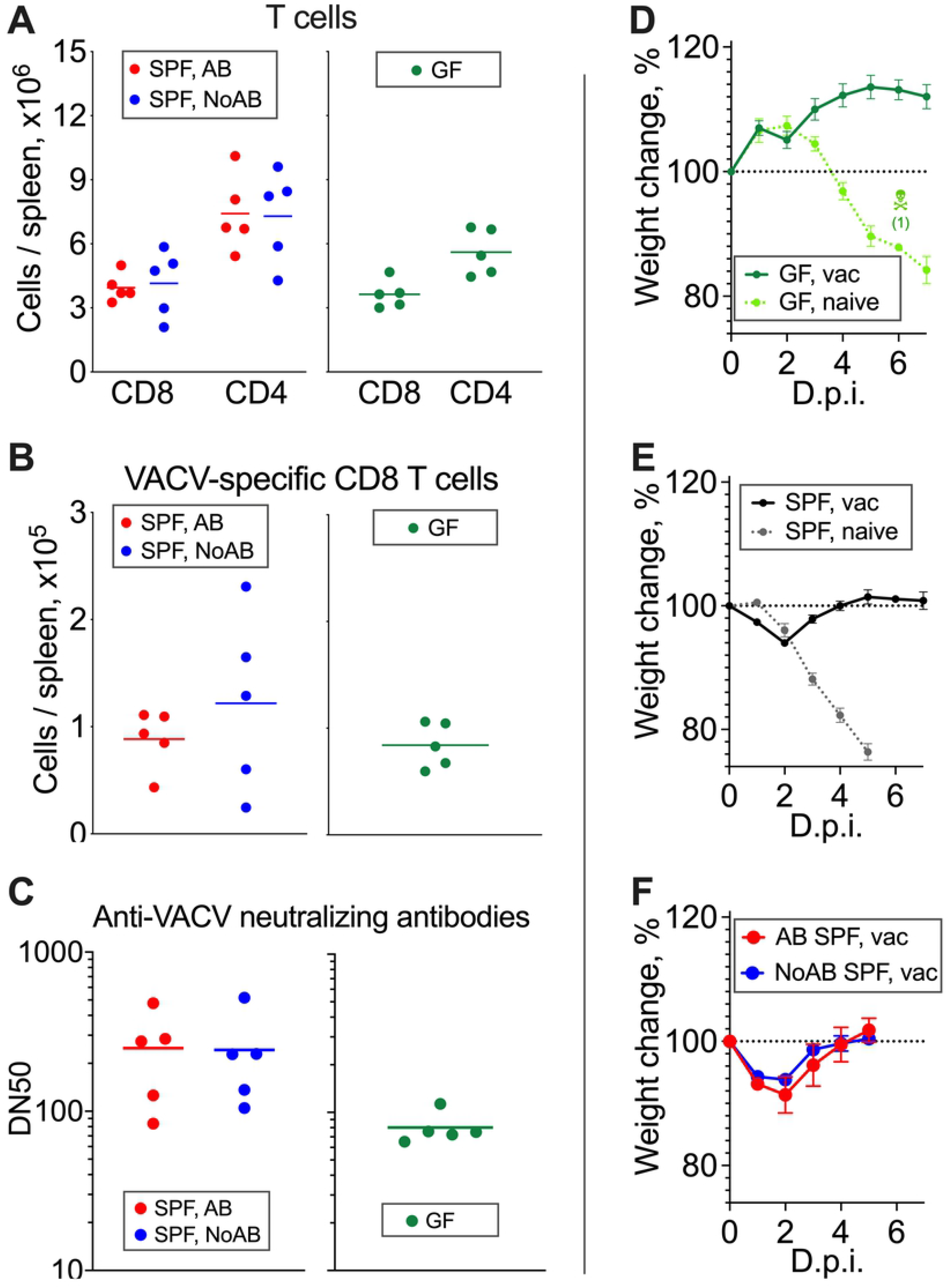
VACV-specific antibody and CD8+ T cell memory response and virus challenge of vaccinated mice. (A-C): SPF or GF mice (*n*=5) were injected i.d. with 10^4^ PFU of VACV. SPF, AB-group received i.p. antibiotic treatment for 10 d from 1 d p.i. SPF, NoAB group received i.p. injections with PBS. Spleens and serum samples were obtained at 34 d p.i. (A) Absolute number of splenic CD8^+^ and CD4^+^ T cells. Bars represent means. (B) Absolute number of VACV-specific splenic CD8^+^ T cells. Bars represent means. (C) VACV-neutralising antibody responses determined by plaque-reduction neutralisation test. IC50, half maximal inhibitory concentration. Bars represent means. All experiments (except with GF mice) were performed twice and representative data from one experiment are shown. The experiments with GF mice were performed once. (D-F): GF and SPF groups of mice (*n*=5 per group) were vaccinated (“vac”) i.d. in both ears with 10^4^ PFU of VACV WR per ear. “Naive” groups (*n*=3-5) were not vaccinated. AB and NoAB groups were treated as in (A-C). One month p.i. groups were challenged i.n. with (D, E) 3 × 10^6^ PFU or (F) 6 × 10^6^ PFU of VACV WR. Data represent the weight of each mouse compared to the weight of the same animal before challenge (d 0). The percentages for each group are means with SEM.

These data do not distinguish between the deficiency of antibody-dependent immunity in GF animals *per se* and the decline in humoral immune responses due to a lack of stimulatory signals from microbiota. We attempted to address this by comparing vaccination of GF and *S. aureus* gnotobiotic mice, which were generated by mono-colonisation of GF animals with *S. aureus* strain NCTC 8325 two weeks prior to the i.d. vaccination with VACV. Unfortunately, mice reconstituted with *S. aureus*, differed from GF controls with lower body mass and reduced subcutaneous and visceral fat, possibly suggest ongoing inflammation after colonisation with *S. aureus*. For these reasons, *S. aureus* colonised animals proved unsuitable for comparison.

## Discussion

Host microbiota influence many physiological and pathological processes throughout life, including pre- and postnatal development, aging, metabolism and immunity (27, 28). Commensal organisms play an important role in the defense against pathogens, not only by preventing microorganisms from colonising epithelial niches, but also by influencing homeostasis, maturation and regulation of the immune system (24, 29). During infection by bacteria or viruses, host microbiota may either promote or diminish pathology (30, 31) and the ability of commensals to influence inflammation via regulation of innate immune responses suggests that microbiota could be manipulated for host health benefit (24, 29, 32).

This study reports a 1000-fold increase in local skin microbiota after i.d. vaccination with VACV (Fig 1). This expanded bacterial load dominates the development of skin lesions following vaccination (Fig 3, S5 Fig) and enhances the innate immune response by increasing leukocyte infiltration and local cytokine/chemokine (Figs 5, 6). Notably, in the absence of microbiota, antibody-mediated immunity is impaired while the antigen-specific CD8^+^ T cell response is preserved (Fig 7).

Multiple factors provide skin barrier function and control microorganisms on the skin surface including acidic pH, high-salt, antimicrobial proteins (defensins, cathelicidins, peptidoglycan recognition proteins, ribonucleases and psoriasin), free fatty acids, ceramide and filaggrin (8, 33, 34). Keratinocytes and sebaceous glands, which synthesise these substances (35, 36), are highly susceptible to VACV infection (37). VACV-induced shut-off of host protein synthesis in infected cells (38) may diminish production of antimicrobial substances and therefore induce changes in local microbiota. Here, VACV infection is shown to induce large changes in skin microbiota, especially later after infection (Fig 2, S2 and S3 Figs), with changing dominance of different bacterial taxa. These dominant taxa varied between infected animals, indicating colonisation by random opportunistic bacteria, a common feature for secondary bacterial infections (39).

The ability of bacterial commensals to rapidly expand has been demonstrated previously; for instance, the doubling time of *S. aureus* in human nasal cavity is 2-4 h (40), which suggests a bacterial increase of 60-2000-fold in 24 h is possible (not accounting for bacterial death and assuming sufficient nutrient availability). The bacterial growth rate in a host organism depends on multiple factors including immune status of the host. During experimental sepsis the speed of bacterial expansion can be more than 100-fold per 24 h, when the immune system loses control of the infection (41). Here we observed up to 100-fold increase of bacterial presence within the first 24 h post VACV-infection (Fig 1C and D), which can be explained by the fact that VACV is a very potent suppressor of the local innate immune response (5). Further bacterial expansion over two weeks post i.d. VACV infection was only 10-fold (Fig 1C and D) possibly slowing down due to nutrient limitations and, after 5 dpi, the activation of the immune system and leukocyte infiltration (Fig 5, Fig 6), which limits bacterial growth and increases their death.

Although increased bacterial, but not viral, load correlated with enhanced lesion size, these increased bacteria were not sufficient for lesion formation, because bacteria increased 1-5 d p.i., before lesions appeared on d 6 (Figs 1C, D and 3A, B). Furthermore, GF mice still developed lesions, although these were delayed and diminished (Fig 3B). Lesion size was also reduced in GF animals following dermal scarification of VACV into the mouse tail (42). VACV infection may disrupt skin integrity due to VACV-induced cytopathic effect (43) or cell motility (44) that enabled bacteria to enter tissue and proliferate. Local immunosuppression induced by VACV proteins that block innate immunity (4, 5) including production of glucocorticoids by a VACV-encoded steroid biosynthetic enzyme (45, 46) might facilitate further bacterial growth. Notably, a VACV strain lacking this enzyme induced smaller dermal lesions (22). In the future, it would be interesting to measure microbiota after infection by this and other VACV mutants lacking specific immunomodulators.

These observations with VACV have similarity to those made with the trypanosome *Leishmania major* in which skin lesions were smaller in GF mice than SPF mice (24). That study also demonstrated the importance of the skin resident immune system in the progression of inflammation (24). Inflammation after exposure to pathogens can be caused and/or aggravated by opportunistic commensals (25). Bacterial invasion provides potent activators of innate immunity that promote cytokine/chemokine production and immune cell recruitment (47). Consistent with this, larger lesions were associated with more neutrophils and T cells (Fig 5B), and increased tissue cytokines and chemokines (Fig 6). Interestingly, depletion of Ly6G^+^ cells results in delayed lesion healing post epicutaneous infection with VACV (37), consistent with the involvement of neutrophils in bacterial clearance and the resolution of inflammation. However, whether bacterial infection *per se* (48) or immune system activation by bacteria (49) induce tissue damage is unknown. During SARS-CoV-2 infection, increased neutrophil counts correlate with severe pathology and hyper-inflammation. Specifically, neutrophils are elevated in the second week from the onset of symptoms and are higher in severe COVID-19 in comparison with moderate disease (50-53). Similarly, the presence of neutrophils predisposes to enhanced respiratory syncytial virus infection (54). In our study, the massive recruitment of immune cells in infected tissue coincided with the appearance of the skin lesion. Therefore, three factors may influence lesion development after i.d. vaccination with VACV: 1) virus-induced tissue damage, 2) tissue destruction by invading bacteria, and 3) immunopathology. Lesion size in this i.d. model is also influenced by the age and strain of mice, and strain of VACV (22).

Bacterial lipopolysaccharides, flagellin, DNA and toxins are used as adjuvants to stimulate innate immunity and vaccine immunogenicity, mainly via enhanced antibody production (17). Consistent with that, the innate immune response to vaccination of GF mice was lower than for genetically-matched SFP animals (Fig 6B), and whereas VACV-specific CD8^+^ T cell memory did not depend on microbiota, the humoral immune response was impaired in GF mice compared with SPF animals (Fig 7A-C). Nonetheless, no reduction of anti-VACV antibodies in antibiotic-treated SPF animals was seen (Fig 7C), despite reduced cytokine/chemokine levels. This maybe because antibiotic treatment does not eliminate bacteria, but limits their growth and spread. Others also reported reduced antibody responses in both GF and antibiotic-treated animals (55, 56). However, these studies used animals pretreated with antibiotic *per os* to induce alterations in the gut microbiome, while we administrated antibiotics i.p. only from 1 d after infection by VACV. Previous studies suggested that the influence of microbiota on T-cell memory may depend on the inoculation route or pathogen (55). Notably, the induction of CD8^+^ T cell memory following i.d. vaccination with VACV is microbiota-independent and not perturbed by the presence of bacteria in the skin, gut or elsewhere (Fig 7).

Although VACV-specific memory CD8^+^ T cells may have functional differences in the absence of microbiota (57), and despite reduced titres of VACV-specific neutralising antibodies, GF animals were protected as well as SPF mice from i.n. challenge with VACV one month (Fig 7D-F). SPF mice that had been treated with antibiotic were also protected as well as control mice at 3 months after vaccination (S7 Fig). This reflects the solid immunity induced by vaccination in all groups and consequential resistance to challenge with a high dose of VACV. These conditions might not be suitable to reveal subtle differences in protection between the groups used. Lower immunising doses or longer duration between vaccination and challenge might show differences. Nonetheless, this study illustrates the influence of skin microbiota for the generation of anti-VACV antibodies and this observation is important for vaccine development.

In conclusion, this study demonstrates that i.d. vaccination with VACV induces substantial local bacterial infection derived from skin microbiota, which function as adjuvant to increase the innate immune response leading to greater skin inflammation. The enhanced bacteria induced formation of larger dermal lesions either by direct bacterial-induced cytotoxicity or immunopathology. Vaccination with VACV generates robust long lasting immune responses that eradicated smallpox from man (1) and can protect mice from lethal challenge (21, 58). As shown here, such protective immunity does not require additional stimuli from microbiota or other adjuvants. However, further activation of the innate immune system provided by skin microbiota might have a beneficial effect on the immunogenicity of other vaccines.

## Materials and Methods

### Animals and study design – ethics statement

C57BL/6 female mice were used in all experiments of this study. Specific pathogen free (SPF) animals were purchased from Charles River and housed under SPF conditions in the Cambridge University Biomedical Services facility. Germ-free (GF) C57BL/6 mice were bred and kept at the University of Bern GF animal facility. All GF mice were confirmed to be microbial-free during breeding and during experiments using culture dependent and independent strategies. Animal experiments in the UK were conducted according to the Animals (Scientific Procedures) Act 1986 under PPL 70/8524 issued by the UK Home Office. Experiments involving GF animals were performed in accordance with Swiss Federal and Cantonal regulations.

SPF mice (7 weeks old) or GF mice (7-9-weeks old) were injected intradermally (i.d.) with 10^4^ plaque-forming units (PFU) of VACV strain Western Reserve (WR) or strain Lister or diluent (0.01% BSA/PBS, mock-control) into both ear pinnae. VACV used for infections was purified from infected cells in sterile conditions by sedimentation through a 36% (w/v) sucrose cushion and subsequently through a 15-40% (w/v) sucrose density gradient. When plated onto sheep blood agar these VACV preparations induced no bacterial growth. VACV titres were measured by plaque assay on BSC-1 cells and stored at -70 °C. The diluted VACV samples used for injections were titrated to confirm the accuracy of the injected dose.

Some groups of animals were injected intraperitoneally (i.p.) with antibiotic or PBS for up to 13 d starting 1 d after i.d. infection with VACV. Ceftriaxone (Rocephin; Roche, Basel, Switzerland), a third-generation cephalosporin antibiotic with a broad spectrum antibacterial activity, was given at 10 mg / mouse / d.

The size of lesions formed after i.d. injection of VACV into the ear pinnae were measured daily with a digital calliper. Whole ear pinnae tissues were collected before and at several times after infection. and spleens and serum samples were obtained one month p.i. to quantify T cell subpopulations and VACV neutralising antibodies. Vaccinated mice and naïve controls were challenged i.n. with 3 × 10^6^ PFU of VACV WR 1 month post vaccination. The body weight of animals was measured daily.

### Flow cytometry

The composition of immune cells in VACV-infected ear tissues was determined by FACS as described (58). Dead cells were excluded by addition of Zombie Fixable Viability dye (S2 Table) for 20 min followed by washing. After pre-incubation of samples with purified rat anti-mouse CD16/CD32 antibody (Mouse BD Fc Block) (BD Biosciences, Cat. # 553141, Franklin Lakes, New Jersey), monoclonal antibodies (mAbs) were added to the cell suspension (S2 Table). The following mAbs were included in the myeloid panel: anti-CD45, Siglec-F, CD11c, CD11b, Ly6C, Ly6G, as well as dump channel markers (CD3, CD5, CD19, NK1.1). The lymphoid panel included CD45, NK1.1, CD3, TCRγ*δ*, CD4, CD8 markers. After final washing steps, cells were resuspended in PBS containing 4% paraformaldehyde (PFA) and were analysed on a BD LSRFortessa (BD Biosciences). Gating strategies are shown in S8 and S9 Figs. FACS analysis were completed using BD FACS Diva (BD Biosciences) and FlowJo (FlowJo, LLC BD, Ashland, Oregon) software.

Splenocyte isolation and staining for FACS analysis were performed as described (58). Subpopulations of splenic T cells were determined by staining with mAbs to CD45, CD3, CD8, CD4 and MHC dextramer H-2Kb/TSYKFESV (S2 Table). Gating strategy is shown in S10 Fig.

### Measurement of cytokines and chemokines in ear tissue

Whole ear pinnae were homogenised in 1.5 ml flat-bottom tubes containing 400 µl of 0.01% BSA/PBS with an OMNI Tissue Homogeniser with plastic hard-tissue probes (OMNI International, Kennesaw, GA, Georgia). The homogenate was then centrifuged at 10,000 rcf for 20 min at 4 °C and the supernatant was collected and stored at -80 °C. The levels of IFNγ, TNFα, IL-1β, IL-4, IL-6, IL-10, IL-33, CCL2, CCL3, CCL4, CCL5, CCL7, CCL20, CXCL1, CXCL2, CXCL5 and CXCL10 were measured using Magnetic Luminex Mouse Premixed Multi-Analyte kits (R&D Systems, Minneapolis, Minnesota) and a Luminex 200 analyser (Luminex Corporation, Austin, Texas).

### Bacterial colony count

Whole ear pinnae homogenates were diluted in 10-fold steps in LB medium and 10 µl of undiluted and diluted samples were pipetted onto sheep blood agar plates (Columbia Agar with sheep blood Plates; Oxoid, Cat. # PB0123A, Thermo Fisher Scientific, Waltham, Massachusetts). After incubation for 18 h at 37 °C, colony-forming units (CFU) were counted. The limit of detection was 40 CFU/ear.

### Viral titres measurement in ear tissues

Whole ears homogenates underwent 3 cycles of freezing-thawing-sonicating to rupture cells and release the virus. Titres of infectious virus were then determined by plaque assay on BSC-1 cell monolayers.

### Histology

Ear pinnae were collected at 8 and 12 d p.i. with VACV strain Western Reserve (WR), fixed in 4% PFA/PBS for 24 h at 4 °C and then stored in 70% ethanol at 4 °C until paraffin embedding. Transverse sections (6 µm) through the middle of the lesion were stained with haematoxylin and eosin (H&E) and were examined under an AxioObserver Z1 microscope with an AxioCam HRc camera (Carl Zeiss AG) using Axiovision software (Carl Zeiss AG).

### Sample preparation for 16S rRNA gene sequencing

Whole ear tissues were collected before infection and at 2, 5, 8 and 12 d post i.d. injection with 10^4^ PFU of VACV WR. Mock-infected samples from mice injected with a sterile diluent (0.01%

BSA/PBS) were obtained at d 8 post injection. Whole ear tissues were homogenised in sterile 1.5 ml flat-bottom tubes containing 400 µl of PowerBead Solution (Part of DNeasy UltraClean Microbial Kit; Qiagen, Cat #12224-250, Hilden, Germany) with an OMNI Tissue Homogeniser with plastic hard-tissue probes (OMNI International). Samples were stored at -80 °C until use. After thawing, 300 µl of homogenates were transferred to PowerBead tubes (Part of DNeasy UltraClean Microbial Kit, Qiagen, Cat #12224-250) and 50 µl of SL solution (Part of DNeasy UltraClean Microbial Kit; Qiagen, Cat #12224-250) was added. The samples then underwent mechanical disruption by rapid agitation with beads using a Fastprep 24-5G bead beater (MP Biomedicals, Fisher Scientific, Irvine, California), 2 cycles of 40 sec at 6 ms^-1^. DNA precipitations were completed with a MasterPure Gram Positive DNA Purification kit (Epicentre, Illumina, Cat # MGP04100, Madison, Wisconsin) according to its instruction manual.

Ear tissue collection and DNA extraction was performed in sterile conditions with single-use instruments and consumables to avoid contamination with bacteria or bacterial DNA. Blank samples containing buffers with no tissue specimens were used as controls for contamination and were treated exactly as ear tissue samples. Blank controls were made for each batch corresponding to the day of tissue collection.

DNA samples were diluted with nuclease free water (Ambion, Cat # AM9935, Austin, Texas) to normalise DNA concentrations to ∼90 ng/µl. The NEXTflex 16S V4 Amplicon-Seq Kit 2.0 (Bio Scientific, PerkinElmer, Waltham, Massachusetts) was used for the bacterial 16S library preparation according to the manufacturer’s instructions with the exception that 150 ng of material was used for amplification with 11 and 22 PCR cycles for the first and second rounds of amplifications, respectively. Kapa Pure beads (KappaBiosystems, Cat # KK8002, Roche) were applied for purification and size selection of DNA with a volumetric ratio of 0.6X-0.8X. The final library was assessed by D1000

ScreenTape assay (Tapestation Agilent; Agilent Technologies, Santa Clara, California) for quality and fragment size, then quantified by Qubit dsDNA High Sensitivity Assay kit (Invitrogen, Thermo Fisher Scientific, Cat. # Q32854) and pooled to 10 nM. The final pool was quantified by qPCR with Kapa Library Quant Kit for Illumina Platforms (KapaBiosystems, Roche, Cat. # KK4824) on an AriaMX Real Time PCR System (Agilent Technologies). Next generation sequencing was performed by paired-end sequencing of the V4 region using MiSeq Reagent Kit version 3, 600-cycle format (Illumina). Library preparation and sequencing were completed by Cambridge Genomic Services. Data have been deposited to the European Nucleotide Archive. Processing and analysis of 16S sequence data is described in S1 Methods.

### Statistical and visual analysis of the microbiome data

#### Heatmap analysis

The heatmap was created using all samples with the R package Heatplus version 2.28.0. (S1 Methods). Taxon was selected if it was present in at least 5 samples with a minimum abundance of 1%. Five clusters were identified by specifying the “cuth” parameter (see S1 Methods). The heatmaps show 50 families from the family taxon datasets, 61 genera from the genus taxon datasets, and 72 species from the species taxon datasets.

#### Principal component analysis (PCA)

For the visualisation of microbial compositional differences between the different sample groups/timepoints we plotted the microbial variances using a multivariate method called Principal Component Analysis (PCA) for the family, genus, and species taxon level using the mixOmics R package version 6.7.1 (59). PCA was conducted including all taxon with a minimum abundance of 0.1%.

### Other statistical analysis

Statistical analysis was performed with SPSS v.25 (IBM, Armonk, New York) and GraphPad Prism v.8.3 (GraphPad Software, San Diego, California). Comparison between two groups of animals were done by Mann-Whitney U-test. The two-way repeated measures (RM) ANOVA test was used for the analysis of time series data. Correlations between parameters were assessed using Spearman correlation coefficient. P values of less than 0.05 were considered significant.

### Data deposit

Sequence data were deposited at the European Nucleotide Archive (ENA) at the European Bioinformatics Institute and will be released upon publication of this article. Accession number PRJEB39345; Study: PRJEB39345 ena-STUDY-Department of Pathology-12-07-2020-12:22:02:769-1371. The other datasets are available on an online open access repository, Figshare. DOI: 10.6084/m9.figshare.14995416

## Acknowledgements

We thank Gillian M. Fraser (Department of Pathology, University of Cambridge) and Stephen D. Bentley (Sanger Institute, Hinxton) for advice on bacteriological aspects of this study.

## Supporting information

**S1 Fig. Morphology of Ly6G+ cells in VACV-infected ear tissue**.

C57BL/6 SPF mice were injected i.d. in the ear pinnae with 10^4^ PFU of VACV WR. Images of sorted neutrophils (Zombie violet^-^, CD3^-^, CD5^-^, CD19^-^, NK1.1^-^, CD11c^-^CD45^+^Siglec-F^-^Ly6G^+^) extracted from ear tissues at 9 d p.i. Red - signal from surface staining with mix of antibodies, blue - DAPI. Bars = 5 µm.

**S2 Fig. Skin microbiome change after i.d. infection with VACV (Family)**.

Ear tissues were collected from SPF mice before (intact) and at 2, 5, 8 and 12 d p.i. with 10^4^ PFU of VACV WR and at d 8 post injection of PBS (mock), *n* = 15 per group per time point. Next generation sequencing was performed on DNA extracted from the ear tissues. (A) Heatmap of bacterial families with hierarchical clustering. Bacterial families with a minimum abundance of 1% in at least five samples was used for creation of the heat map. Five clusters are colour coded with red, blue, green, violet and orange. (B) Results of Principal Component Analysis (PCA) of ear tissue microbiome of family taxon. PCA was conducted including all taxon with a minimum abundance of 0.1%. Ellipses represent a 40% confidence interval around the cluster centroid. “D” – day post i.d. injection.

**S3 Fig. Skin microbiome change after i.d. infection with VACV (Species)**.

Ear tissues were collected from SPF mice before (intact) and at 2, 5, 8 and 12 d p.i. with 10^4^ PFU of VACV WR and at d 8 after injection of PBS (mock), *n* = 15 per group per time point. Next generation sequencing was performed on DNA extracted from the ear tissues. (A) Results of Principal Component Analysis (PCA) of ear tissue microbiome of specie taxon. PCA was conducted including all taxon with a minimum abundance of 0.1%. Ellipses represent a 40% confidence interval around the cluster centroid. (B) Relative abundance of most prevalent taxa of ear tissue bacterial species. (C) Heatmap of bacterial species with hierarchical clustering. Bacterial species with a minimum abundance of 1% in at least five samples was used for creation of the heat map. Five clusters are colour coded with red, blue, green, violet and orange. “D” – day post i.d. injection.

**S4 Fig. Dermal lesion sizes after antibiotic treatment of animals infected with VACV strain Lister**. Groups of C57BL/6 SPF mice (*n*=5 per group) were injected i.d. with 10^6^ PFU of VACV strain Lister. Group AB were injected i.p. with antibiotic ceftriaxone daily from d 1 to 11 p.i. Group NoAB received i.p. injections with PBS. Data show the mean lesion size +/- SEM. Statistical analysis by two-way RM ANOVA test, p=0.018.

**S5 Fig. Dermal lesion sizes after antibiotic cream treatment of animals infected with VACV**.

Groups of C57BL/6 SPF mice (*n*=3-5 per group) were injected i.d. with 10^4^ PFU of VACV strain WR. The first group received topical administration with 3% tetracycline cream on infected ears daily from d 5 to 21 p.i. The second group received topical administration with 4% erythromycin cream on infected ears daily from d 5 to 21 p.i. Group NoAB received topical application of neat vaseline. Data show the mean lesion size +/- SEM. Statistical analysis by two-way RM ANOVA test.

**S6 Fig. Antibiotic treatment does not influence leukocyte composition of blood, spleen and bone marrow**.

C57BL/6 SPF mice received i.p. injections with antibiotic (ceftriaxone) or PBS for 10 d, then numbers of different subpopulations of myeloid and lymphoid cells were measured in blood (A), spleen (B) and bone marrow (C) (n=5 per group). DC & MФ, dendritic cells and macrophages; Mon, monocytes, TCRγδ, TCRγδ^+^ T cells; TCRαβ CD4, TCRγδ^-^CD4^+^ T cells; TCRαβ CD8, TCRγδ^-^CD8^+^ T cells. Box plots are shown.

**S7 Fig. VACV-specific antibody and CD8+ T cell memory response and virus challenge of mice 3.5 months post vaccination**.

(A-C): SPF or GF mice (*n*=5) were injected i.d. with 10^4^ PFU of VACV. SPF, AB-group received i.p. antibiotic treatment for 10 d from 1 d p.i. SPF, NoAB group received i.p. injections with PBS. Spleens and serum samples were obtained at 3.5 months p.i. (A) Absolute number of splenic CD8^+^ and CD4^+^ T cells. Bars represent means. (B) Absolute number of VACV-specific splenic CD8^+^ T cells. Bars represent means. (C) VACV-neutralising antibody responses determined by plaque-reduction neutralisation test. IC50, half maximal inhibitory concentration. Bars represent means. (D): SPF mice were vaccinated i.d. in both ears with 10^4^ PFU of VACV WR per ear. AB and NoAB groups (*n*=5 per group) were treated as in (A-C). At 3.5 months p.i. groups were challenged i.n. with 10^7^ PFU of VACV WR. Data represent the weight of each mouse compared to the weight of the same animal before challenge (d 0). The percentages for each group are means with SEM.

**S8 Fig. Gating strategy for flow cytometry of myeloid lineage cells present in ear tissue 7 d after i.d. injection of SPF mice with 10**^**4**^ **PFU of VACV WR**.

Cells were gated on their ability to scatter light. Doublets were excluded using a FSC-A versus FSC-H plot. The myeloid gate included CD45^+^Zombie dye^-^CD3^-^CD5^-^CD19^-^NK1.1^-^ cells. Further myeloid cell subpopulations were classified as follows: Eosinophils: CD45^+^CD3^-^CD5^-^CD19^-^NK1.1^-^CD11c^-^Siglec-F^+^; DC and MФ (dendritic cells and macrophages): CD45^+^CD3^-^CD5^-^ CD19^-^NK1.1^-^Siglec-F^-^CD11c^+^; Neutrophils: CD45^+^CD3^-^CD5^-^CD19^-^NK1.1^-^CD11c^-^Siglec-F^-^ Ly6G^+^; Ly6C^+^Mon (inflammatory monocytes): CD45^+^CD3^-^CD5^-^CD19^-^NK1.1^-^CD11c^-^Siglec-F^-^Ly6G^-^CD11b^+^Ly6C^+^; Ly6C^-^Mon (residential monocytes): CD45^+^CD3^-^CD5^-^CD19^-^NK1.1^-^CD11c^-^Siglec-F^-^Ly6G^-^CD11b^+^Ly6C^-^.

**S9 Fig. Gating strategy for flow cytometry of lymphoid lineage cells present in ear tissue 7 d after i.d. injection of SPF mice with 10**^**4**^ **PFU of VACV WR**.

Cells were gated on their ability to scatter light. Doublets were excluded using a FSC-A versus FSC-H plot. Then, hemopoietic cells were gated as CD45^+^Zombie dye^-^ cells. Further lymphoid subpopulations were classified as follows: NK cells: CD45^+^CD3^-^NK1.1^+^; TCRγδ T cells: CD45^+^CD3^+^NK1.1^-^TCRγδ^+^; CD4 T cells: CD45^+^CD3^+^NK1.1^-^TCRγδ^-^CD8^-^CD4^+^; CD8 T cells: CD45^+^CD3^+^NK1.1^-^TCRγδ^-^CD4^-^CD8^+^.

**S10 Fig. Gating strategy for flow cytometry of splenic CD8**^**+**^ **and CD4**^**+**^ **T cells isolated one month after i.d. injection of ear pinnae with 10**^**4**^ **PFU of VACV WR, or from mock-infected animals**. Cells were gated on their ability to scatter light. Doublets were excluded using a FSC-A versus FSC-H plot. T cells were classified as CD45^+^Zombie dye^-^CD3^+^ cells and then divided into CD8^+^ and CD4^+^ T subsets and with further gating of VACV-specific CD8^+^ T lymphocytes (CD45^+^CD3^+^CD4^-^CD8^+^Dextramer^+^).

**S1 Table. Variation in bacterial composition (beta diversity) among groups of samples at different times p.i. as well as mock and intact ear specimens by PERMANOVA test with and without Bonferroni correction**.

**S2 Table. Monoclonal antibodies and dyes used for staining cells prior to analysis by flow cytometry**.

